# Missing genomic resources for the next generation of environmental risk assessment

**DOI:** 10.1101/2023.10.11.561851

**Authors:** Marc Sven Roell, Mark-Christoph Ott, Magdalena M. Mair, Tobias Pamminger

## Abstract

Environmental risk assessment traditionally relies on a wide range of *in vivo* testing to assess the potential hazard of chemicals in the environment. These tests are often time-consuming, costly and can cause test organisms’ suffering. Recent developments of reliable low-cost alternatives, both *in vivo*- and *in silico*-based, opened the door to reconsider current toxicity assessment. However, many of these new approach methodologies (NAMs) rely on high quality annotated genomes for surrogate species of the regulatory risk assessment. Currently, lacking genomic information slows the process of NAM development. Here, we present a phylogenetically resolved overview of missing genomic resources for surrogate species within regulatory ecotoxicological risk assessment. We call for an organized and systematic effort within the (regulatory) ecotoxicological community to provide these missing genomic resources. Further, we discuss the potential of a standardized genomic surrogate species landscape to enable a robust and non-animal reliant ecotoxicological risk assessment in the systems ecotoxicology era.

**Synopsis Statement:** We identify missing genomic resources needed for the development and regulatory acceptance of new approach methodologies in environmental risk assessment.

## Introduction

Traditional dose-response toxicity tests are a fundamental building block of regulatory ecotoxicology and the preferred method for *in vivo* quantification of apical hazards ^1^. However, the dogma of traditional dose-response toxicity tests as “gold standard” is questioned due to the neglected consideration of biological variability, lack of assay robustness and the suffering and resulting sacrifice of many organisms involved in such tests. In addition, such classic *in vivo* assays are not designed to elucidate the mechanistic bases of the observed adverse effects which often makes their interpretation challenging ^2, 3^. A shift in political, societal, and scientific perception promotes a new generation risk assessment (NGRA) based on the mechanistic understanding of adverse effects. Fundamental within the evolving topic of NGRA in many regulatory areas are new approach methods (NAMs) ^4,5^. Historically, NAMs were intended for mechanistic hazard characterization in the 3R vertebrate domain ^6^. Recently, these approaches have been extended beyond their original narrow phylogenetic scope to a broad range of organisms ^7^. NAMs deviate from traditional dose-response toxicity tests. They comprise a heterogeneous collection of approaches that include a variety of *in vitro* experimental setups (e.g., cell lines and embryos), experimental readouts (OMICs -RNAseq, proteomics, metabolomics), and in silico methods (e.g., Bayesian networks and machine learning applications). Further, NAMs offer potential for implementation within high-throughput applications to characterize adverse effects at scale (for a detailed view on NAMs see ^7^).

Despite their diversity, many NAMs rely on sub-organismic measurement data that require high-quality genomic information throughout the design, validation, and interpretation phases (e.g., sequencing depth and functional annotation). Prominent examples use robust hazard-associated biomarkers to predict adverse effects in a range of study organisms (e.g. EcoTox Chip) ^8^ or characterize predictive quantitative adverse effects (qAOPs) ^9^.

The clear benefits of available high-quality genomic resources for standard ecotoxicological surrogate species remain a promise of the ecotoxicogenomics revolution and have yet to be fulfilled ^10^. Among other reasons, the prohibitive high costs of genome sequencing and low collaborative efforts to establish a community-wide genome atlas have likely contributed to a fractionated genomic landscape across ecotoxicological surrogate species.

The dramatic decrease in sequencing costs around 2007 enabled a broad applicability for genome sequencing and promoted the development of computational analysis tools. Now, global consortia generate genomic biodiversity biobanks via large-scale high quality genome sequencing and might be an opportunity to develop a high-quality genome atlas for regulatory relevant surrogate species ^11, 12, 13^.

To support the development of a harmonized open-access genome atlas as a community-wide genomic resource for (regulatory) ecotoxicology, we have compiled a list of currently available standard test species in EU and USA regulatory frameworks (i.e., species mentioned in standardized test guidelines) and combined it with information on phylogeny, habitat, genome resource availability from NCBI and genome quality. We will use this data to provide an overview of the availability and distribution of genomic resources. We hope that a field-wide push to close genomic resource gaps will accelerate the transition towards NGRA and help to facilitate the full transition of ecotoxicology into the 21^st^ century.

## Material & Methods

In spring 2022, we screened standardized ecotoxicological test guidelines (i.e. OECD, OCSPP and ASTM) for recommended surrogate species named at the species or genus level. Communities and test organism where the regulatory relevant species could not be unambiguously assigned were excluded from the analysis (e g. microbial community was excluded while *Bufo bufo* was considered as representative species for *Bufo sp*.). In total, seven entities were excluded from the analysis (Gossypium spp., microbial community, Procambarus sp., Protozoan community, Rhizobium sp., Uca sp.). For all organisms we recorded their habitat (freshwater, salt water or terrestrial) and the taxonomic group they belong to (invertebrates, vertebrates plants and algae; data taken from EnviroTox ^14^). Using this organism list, we searched for assembled genomes deposited in the NCBI database (search date June 2022) using the genome search function. For each species we recorded the presence or absence of an assembled genome and the link to this resource. Assembled genomes include, as defined by the NCBI, whole genome sequences, genomic maps, chromosomes, assemblies, and annotations (ESM Table S1).

The genome sequence quality in terms of coverage and assembly was evaluated in a stepwise approach for each species. First, we assessed if the available genome assemblies covered the chromosomal level, a key quality criterion ^15^. Second, following recent guidance ^15^, quality metrics for each genome were assessed. We defined genomes as high-quality, if a BUSCO score (Benchmarking Universal Single-Copy Orthologs) above 90% was given and if the ratio of contigs to chromosome pairs (CC ratio) was below 1000 ^15^. For organisms named at the genus level in guidance documents, the genome quality of a representative species was evaluated.

For visualization, we constructed a consensus phylogeny for the available regulatory surrogate species based on data obtained from the Open Tree of Life Project using the R package *rotl* ^16^.

To this end, organism names from guidance documents were matched with species names in the open tree taxonomy (synthesis release v14.7) ^17^. Matches were checked carefully for duplicate names, duplicate database IDs (ott numbers), assignments to correct taxonomic domains, and synonymous species names. Entries with species names of unclear taxonomic status were checked and adjusted based on manual searches on external taxonomic databases (World Register of Marine Species ^18^, Catalogue of Life ^19^). For unresolved species names, genus nodes or nodes of most recent common ancestors were chosen for tree construction, if possible. Species for which an assignment of a unique number was not possible due to unknown phylogenetic status were not considered. Synthetic trees were visualized together with information on habitat, availability of assembled genomes and their respective quality with *ggtree* ^20^ and *ggplot2* ^21^. All analyses were done in R version 4.2.1 ^22^.

All data and R code will be provided on acceptance. The raw data files are available in the ESM.

## Results

In total we found 263 mentioned biological entities in the screened regulatory guidance documents (Figure 1). All entities, where the regulatory relevant species are not unambiguously assigned,were excluded from the analysis (e g. microbial community was excluded while *Bufo bufo* was considered as representative species for *Bufo sp*.) In total, 104 out of 256 (40.6% of all species) species-specific genomic resources were available at NCBI (ESM Table S1). 75 of these genomes were resolved on a chromosome level, a key quality criterion ^14^ (ESM Table S2). Based on the quality metrics defined previously ^15^, 17 genomes were not categorized as high-quality due to either a BUSCO score below 90% (*Gadus morhua, Mytilus edulis, Secale cereale*) or a calculated CC ratio above 1000 (14 additional species, ESM Figure S1). 58 genomes remained as high-quality categorized genomes (22.6% of all species, see ESM Table S2).

**Figure 1.**
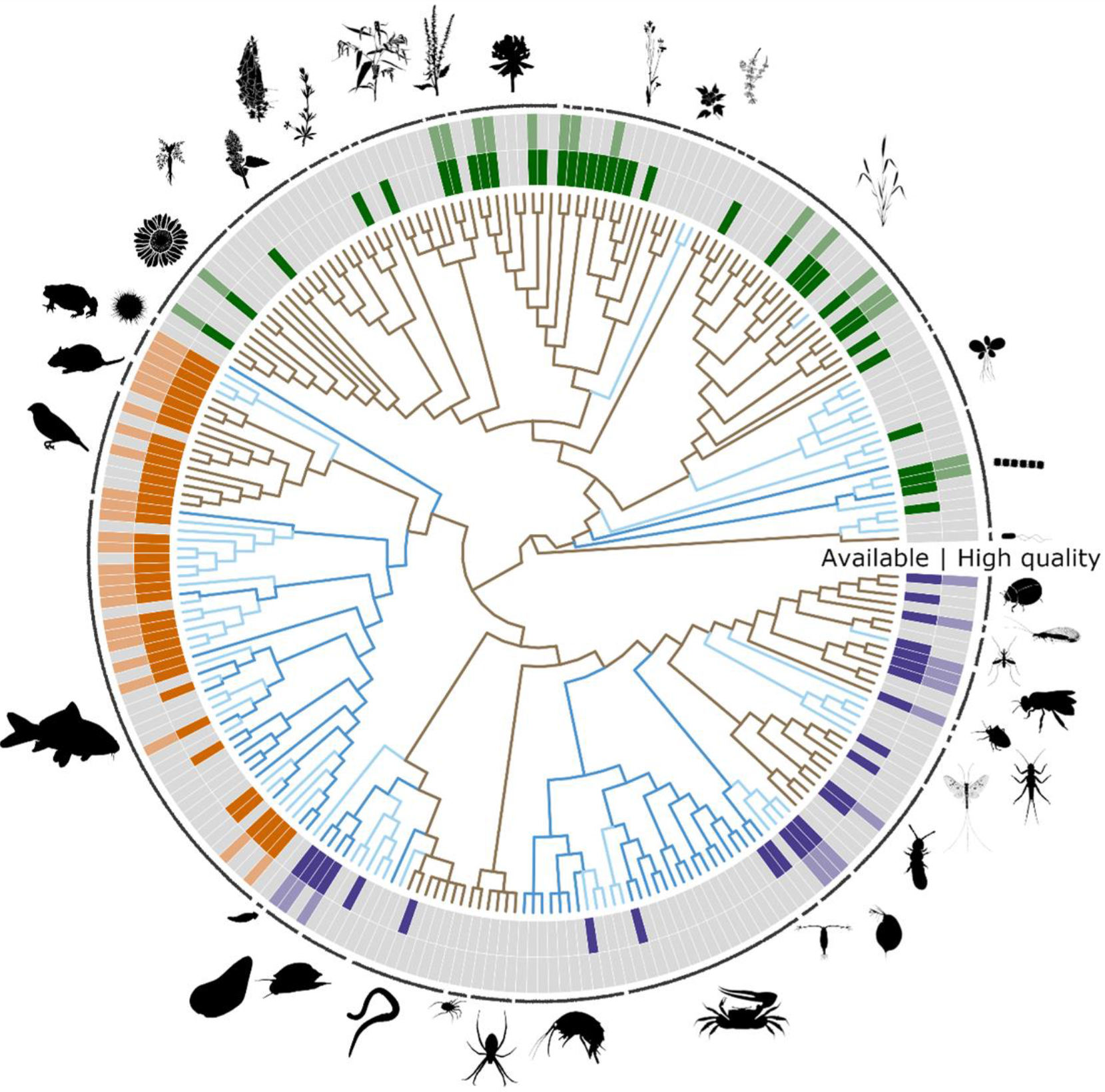
Availability of genomic resources (inner circle) and their quality (outer circle) for test species mentioned in standardized ecotoxicological test guidance documents used for regulatory risk assessments. Within the phylogeny the habitat of the species is indicated by color (brown = terrestrial, dark blue = salt water, light blue = fresh water).

Six additional species are currently sequenced by large sequencing initiatives (Darwin Tree of Life project ^13^, i5k ^23^, Aqua-FAANG ^24^, the International Weed Genome Consortium ^25^) likely resulting in high quality genomes which were not considered in the current analysis (see ESM Table S3). We found that both the available and high-quality genomic resources have unequal coverage among the large taxonomic groupings with vertebrates having the highest total number and percent coverage, followed by plants and invertebrates (Table1). An individual analysis on the phylogenetic genomic resource availability for plants and algae, invertebrates and vertebrates are shown in the ESM (Figures S2-S4).

**Table 1.**
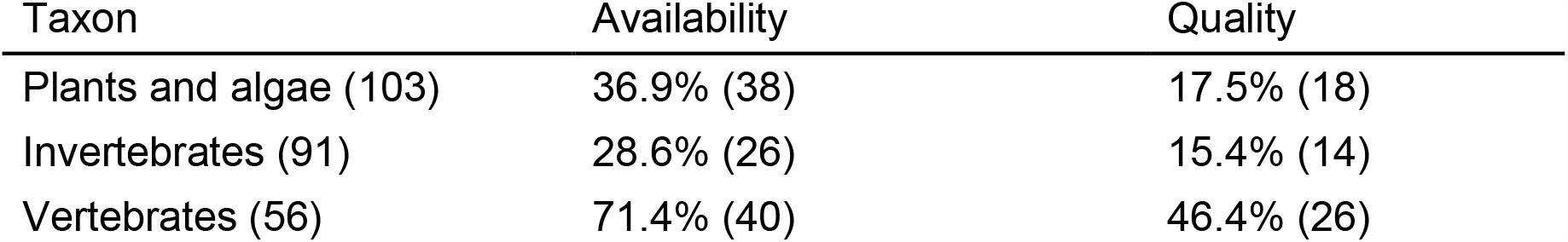
Percentage of species, for which genomic resources are available and proportion of species for which genomic resources are available in good quality. Total Numbers shown in brackets.

| Taxon | Availability | Quality |
| --- | --- | --- |
| Plants and algae (103) | 36.9% (38) | 17.5% (18) |
| Invertebrates (91) | 28.6% (26) | 15.4% (14) |
| Vertebrates (56) | 71.4% (40) | 46.4% (26) |

## Discussion

We systematically assessed both the availability and quality of the genomic resources for surrogate test species used in regulatory ecotoxicology. We found that overall, less than 23% of the investigated surrogate species are covered by genomes of high quality.

Since the beginning of the genomic era in the early 2000s, the potential application of toxicogenomics in (eco)-toxicology research has been obvious and it was considered a logical step towards predictive ecotoxicology ^10^. Sequencing genomes was a time-consuming and prohibitively costly endeavor at the time resulting in sparse and biased coverage of relevant model species. Recent drops in costs and the advent of the third-generation long-range DNA sequencing and mapping technologies are now creating a renaissance in high-quality genome sequencing. Unlike second-generation sequencing, which produces short reads a few hundred base-pairs long, third-generation single-molecule technologies generate over 10,000 bp reads or map over 100,000 bp molecules. This results in major improvements of the quality of genome assemblies, allows to resolve the most challenging genomic regions and to generate telomere-to-telomere assemblies of whole chromosomes ^265^. Genome sequencing projects like the Earth Biogenome ^11^ or Darwin Tree of Life ^13^ are generating reference genomes for many known species as a resource for future bioscience and environmental stewardship. Given these advances, it is now feasible to correct the historic coverage and a high-quality open access ecotoxicology-relevant genome atlas can be generated. We consider such a genome atlas a key puzzle piece for NAM development. At its core, a genome atlas provides robust and consistent genome annotations for a set of species. In an ecotoxicological context, such an atlas would expand the value sequence alignment to predict across species susceptibility (SeqAPASS ^27^) and facilitate the description and characterization of adverse outcome pathways at a molecular level. Based on a detailed molecular understanding of toxicity generating processes including conserved outcome pathways, detoxification mechanisms and physiological responses, toxicity extrapolations to multiple species beyond closely related ones will become possible ^28, 29^.

Beyond its central role for NAM development and -validation in regulatory ecotoxicology, we are convinced that a high-quality genome atlas will provide a robust foundation for a holistic understanding of the effects of chemical stressors from their molecular initiating events up to observed adverse effects at an organismic level. Recent experimental and computational advances have paved the way towards such an integrated view of complex biological processes and systems (from molecules to organisms), known as systems biology ^30, 31^. The interdisciplinary field of systems biology aims to understand biological systems, rather than studying individual components in isolation. Such reasoning is directly applicable to chemical induced stress responses and can support a comprehensive understanding of the impacts of toxic substances on organisms, populations and finally ecosystems ^30, 31^. Within a systems ecotoxicology framework, we envision the development of a multi-tool NAM based approach that covers OMICs (genomics, single-cell transcriptomics, proteomics, metabolomics), high-density biological data cellular- or bioassays) and *in silico* methods (toxicokinetic-toxicodynamic [TK-TD] and dynamic energy budget [DEB] modelling) to explain synthetic chemical toxicity mechanistically and systematically in potential off-target species. Supported by such a multifactorial data integration system ^32^, it will become possible to move beyond simplistic toxicity proxies such as LC/EC50 values and to understand adverse effects caused by chemical stressors in a mechanistically informed manner. Such a framework will hopefully ensure more robust predictions of adverse outcomes across levels of biological organization and strengthen more sustainable and ethical solutions for the urgent questions related to food production and adaptation to global change societies face today.

## Supporting information

ESM

