## Supplementary material for "Missing genomic resources for the next generation of environmental risk assessment": ESM

### Extended Supporting Material (ESM):

Table S1: List of recommended surrogate species in standardized ecotoxicological test guidelines. Test guidelines in which the respective surrogate species are mentioned are listed in the reference column. Link to the genome based on the NCBI search is provided as well.

Table S2: List of genomes resolved at chromosome levelfor surrogate species in standardized ecotoxicological test guidelines. Contig/Chromosome ratio = CC ratio [15]. Chromosome counts in italics derived from the Animal Chromosome Count Database [33]

Table S3: Additional species sequenced in large sequencing consortia.

| Genome Sequencing project | Organism |
|---|---|
| Darwin Tree of Life project | Enchytraeus crypticus |
| | Lolium perenne |
| International Weed Genome Consortium | Bromus tectorum |
| | Avena fatua |
| | Echinochloa crusgalli |
| | Cyperus rotundus |

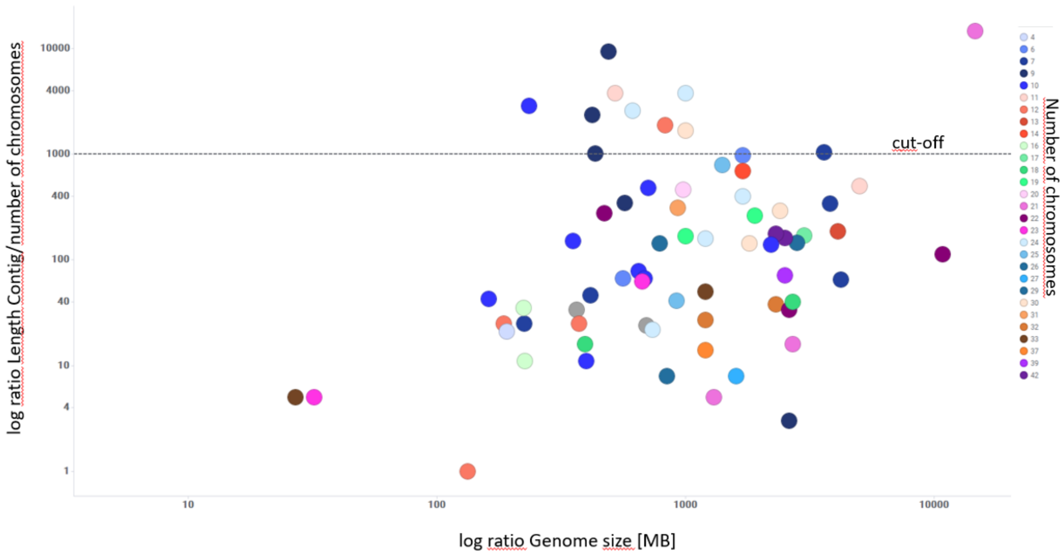


Fig S1. Log ratio of contig length/number of chromosomes (CC) versus log genome sizes in megabases (MB).

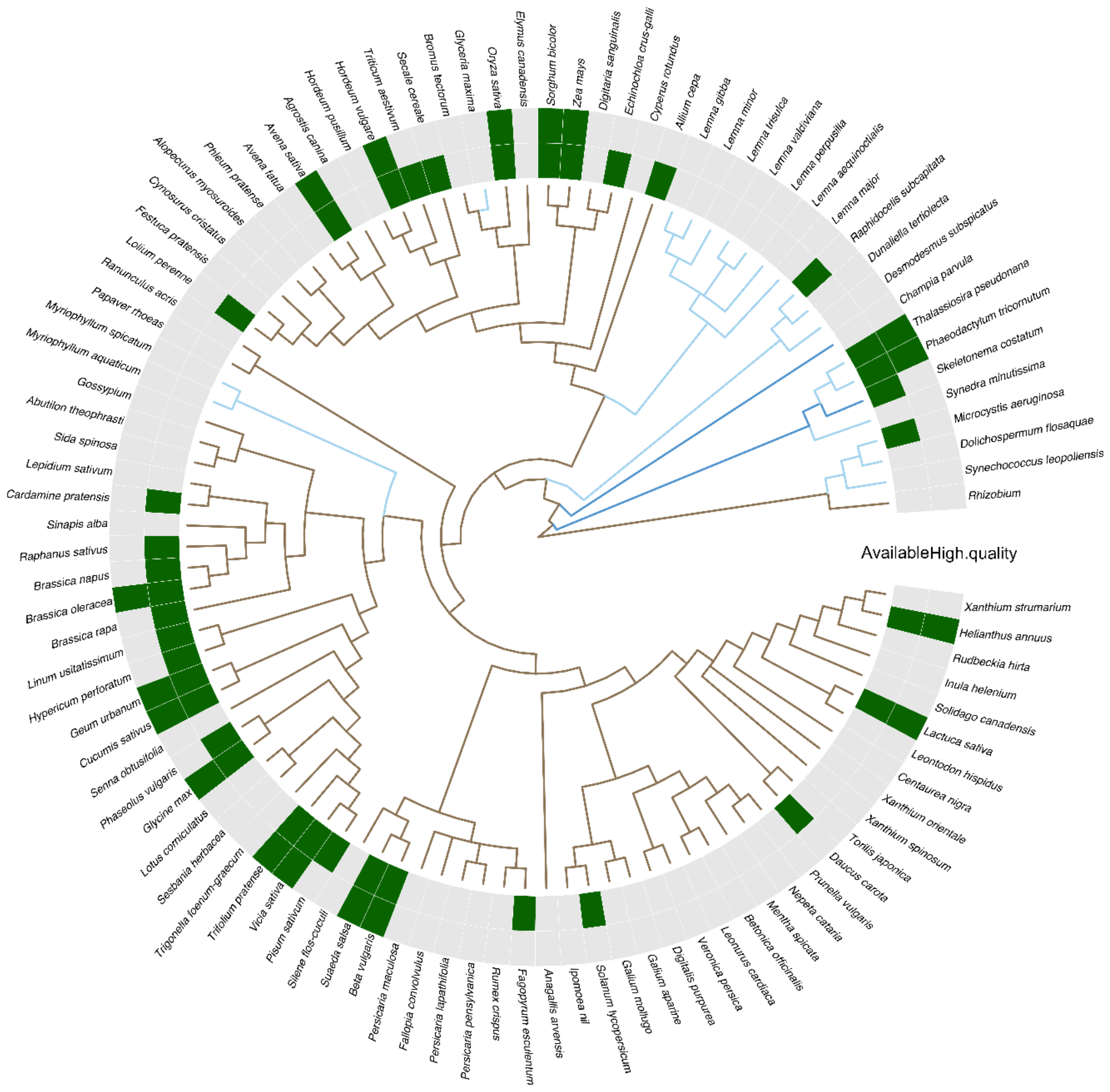


Figure S2. Availability of genomic resources (inner circle) and their quality (outer circle) for plants and algae mentioned in standardized ecotoxicological test guidance documents used for regulatory risk assessments. Within the phylogeny the habitat of the species is indicated by color (brown = terrestrial, dark blue = salt water, light blue = fresh water).

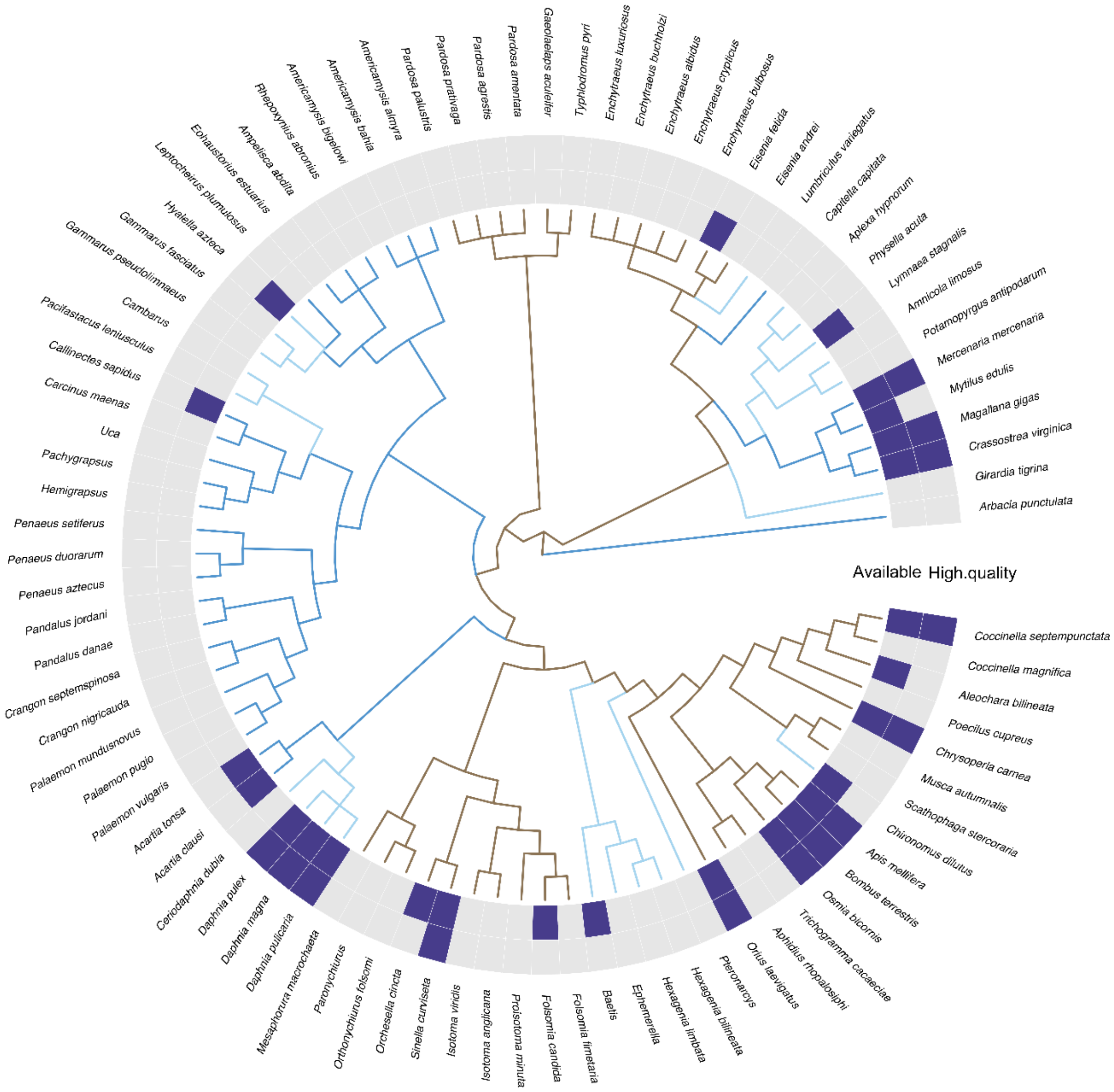


Figure S3. Availability of genomic resources (inner circle) and their quality (outer circle) for invertebrates mentioned in standardized ecotoxicological test guidance documents used for regulatory risk assessments. Within the phylogeny the habitat of the species is indicated by color (brown = terrestrial, dark blue = salt water, light blue = fresh water).

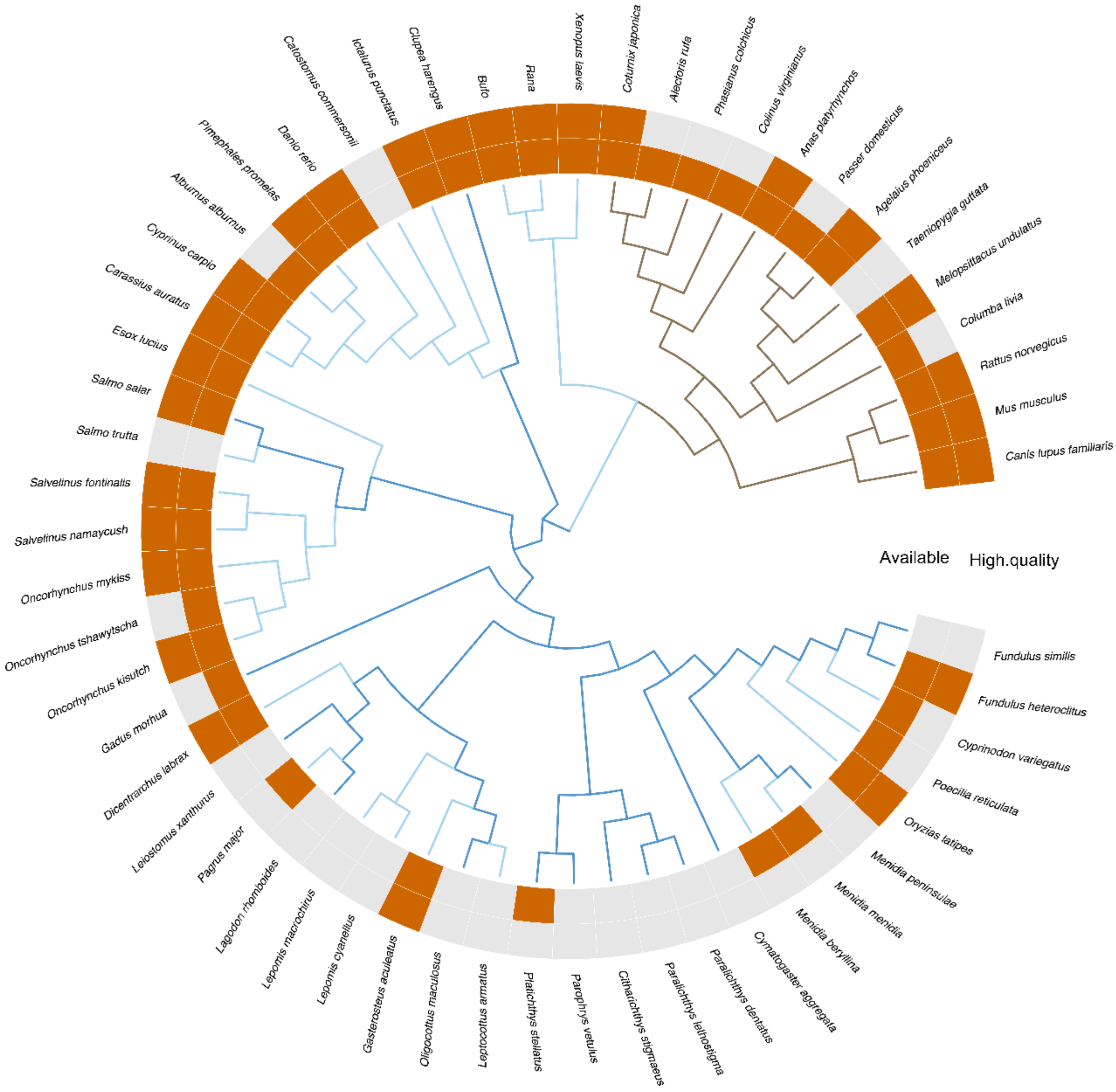


Figure S4. Availability of genomic resources (inner circle) and their quality (outer circle) for vertebrates mentioned in standardized ecotoxicological test guidance documents used for regulatory risk assessments. Within the phylogeny the habitat of the species is indicated by color (brown = terrestrial, dark blue = salt water, light blue = fresh water).
